# Utilizing mentorship education to promote a culturally responsive research training environment in the biomedical sciences

**DOI:** 10.1101/2023.08.25.554846

**Authors:** Sarah Suiter, Angela Byars-Winston, Fátima Sancheznieto, Christine Pfund, Linda Sealy

## Abstract

There is an urgent and compelling need for systemic change to achieve diversity and inclusion goals in the biomedical sciences. Since faculty hold great influence in shaping research training environments, faculty development is a key aspect in building institutional capacity to create climates in which persons excluded because of their ethnicity or race (PEERs) can succeed. In this paper, we present a mixed methods case study of one institution’s efforts to improve mentorship of PEER doctoral students as a strategy to improve graduate trainees’ experiences, and as a strategy to positively affect institutional climate with respect to racial and ethnic diversity. We found evidence that our culturally responsive mentorship education increased faculty mentors’ self-reported mentorship competencies, though their valuing of culturally responsive mentoring behaviors were comparatively higher than their confidence to enact those behaviors. PEER doctoral students reported that they noticed mentors’ efforts to address cultural diversity matters and identified some guidance for how to approach such topics. We discuss future directions and implications for using mentorship education to activate systemic change toward inclusive research training environments and promoting the value of mentorship within institutions.

## Introduction

Lack of racial and ethnic diversity in science, technology, engineering, and math (STEM) fields is a well-documented problem [1–3], one that begins in undergraduate education and worsens as students advance through their academic pathways and into STEM careers [4]. The National Center for Science and Engineering Statistics found that persons excluded from science and higher education because of their ethnicity or race (PEERs) enter college with similar or higher rates of intention to pursue a STEM degree, but their representation diminishes at each stage of higher education [2, 5–6]. For example, in 2019, PEERs constituted 30% of the US population, but earned 18% of STEM bachelor’s degrees and 11% of STEM PhDs [2]. Underrepresentation persists in the scientific workforce, where 13% of the STEM workforce and only 4% of faculty at research institutions identify with a minoritized racial or ethnic group [3]. These disparities are consequential for STEM fields themselves, as racial diversity is associated with increased innovation and other forms of scientific excellence [7]. They are also evidence of long-standing injustices that higher education has an obligation to address [5, 8–9].

The last few decades have seen concerted efforts to address the lack of racial diversity in STEM, with program and policy initiatives aimed at a variety of levers of change, including student-level interventions, institution-level interventions, and interventions that target relationships with peers and mentors [1]. Initially, most interventions were focused on PEERs themselves, and sought to address issues such as lack of identification with STEM disciplines, lack of self-efficacy and preparation, and increased access to resources and support services [10]. Although some of these interventions demonstrated success [11], those that failed to contextualize student-level issues (e.g., lack of preparation) within a broader framework of educational injustice often failed to address the true needs of students in systems and institutions not designed for their success [12, 13].

In this paper, we present a case study of one institution’s efforts to improve mentorship of PEER doctoral students as a strategy to improve graduate trainees’ experiences, and as a strategy to positively affect institutional climate with respect to racial and ethnic diversity. The culturally responsive mentorship education program, described in a later section, was one institutional-level intervention among many implemented by the university to affect organizational learning needed for systemic change. Based on the lack of racial and ethnic diversity in STEM fields noted in the opening paragraph and in other publications [14], this culturally responsive mentorship education program focuses largely on racial and ethnic diversity matters in research mentoring relationships. Through this case study, we explored faculty experiences with and attitudes toward mentorship, their intentions to implement mentoring changes post-workshop, and student mentee perspectives on faculty mentoring after the workshop. Where specific authors contributed uniquely to the research design or data analyses, we note this contribution with their initials.

### Mentorship Education as a Systemic Intervention

There is an urgent and compelling need for systemic change to achieve diversity and inclusion goals in STEM [15] because it is the research training systems that need to be fixed, not the students. Since faculty hold great influence in shaping research training environments, faculty development is a key aspect in building institutional capacity to create climates in which students from more demographically diverse backgrounds can succeed. Hurtado et al. [17] specified an organizational learning model to illustrate how systemic change at the graduate department level in STEM disciplines may be catalyzed by faculty who participate in institutional-level faculty development interventions. Their model begins with science faculty gaining new knowledge about the experiences of trainees, especially those from PEER groups, followed by interventions that can change faculty mindsets and behaviors. In turn, diffusion of new knowledge can lead to critical questioning of existing departmental and campus policies and practices that can be hostile to PEER students. Ultimately, critical questioning of the institutional status quo research training practices necessitates greater faculty buy-in to overcome resistance due to deeply embedded elements of culture and power dynamics that must be confronted in order to sustain new inclusive practices aimed at student success. Hurtado’s model of organizational learning helps describe how institutional interventions through faculty development, like mentorship education, can promote broader change to improve inclusive graduate training in STEM.

The NASEM report, *The Science of Effective Mentorship in STEMM* defines mentorship as a “professional, working alliance in which individuals work together over time to support the personal and professional growth, development, and success of the relational partners through the provision of career and psychosocial support” (p.37) [8]. As highlighted in the report, mentoring relationships play a pivotal role in student academic outcomes such as degree completion, academic productivity, and job acquisition, as well as psychosocial outcomes like sense of belonging, anxiety, and depression [8, 16]. Effective mentors help students access resources and navigate institutional systems [6], create positive micro-climates within their own laboratories and research teams [17, 18], and play a direct role in socializing students to STEM fields and careers [13]. The impact of mentorship is even more pronounced for PEERs studying at predominantly white institutions [19]. Most faculty, however, do not receive formal mentorship education, and many institutions do not provide the types of requirements and/or incentives that communicate to faculty and students that mentorship is something they value [16]. Nevertheless, mentorship education has the potential to improve effective mentorship knowledge and practices, promote the value of mentorship within institutions and activate systemic change toward an inclusive science [14, 20, 24].

Our case study examines mentorship education efforts in the biomedical sciences at Vanderbilt University and is distinct from previous studies on mentoring interventions in STEM in at least two ways. First, unlike many mentor training programs in which participants are selected for participation based on their experience with, commitment to, and talent for mentoring *prior* to the training, this training was open to all faculty in specific departments and programs at various time points. As a result, we are able to explore variations in motivation for faculty participation, experience with mentoring PEER students, and perceptions of mentorship in a more generalized group of faculty located in one institution. Second, the use of qualitative data collection strategies to gain the perspectives of mentees themselves provides rich information with respect to student experiences of mentorship as their mentors implement improvements to their mentoring practices.

## Case Study

The Basic Sciences of the Vanderbilt School of Medicine and the research enterprise of Vanderbilt University Medical Center were well-positioned to implement mentorship education as a strategy to catalyze institutional change toward inclusive excellence. As a consequence of a 20+ year commitment to diversity in graduate education, on average 25% of doctoral trainees in biomedical research training programs are from PEER groups. Overall, Vanderbilt has awarded 246 doctoral degrees in biomedical research to students from PEER groups since 1997, and in 2016 they were the top producer of African American doctorates in the biomedical sciences in the country. The success in biomedical graduate student diversity at Vanderbilt has come about in large measure through the early adoption of holistic admissions processes. Beginning in 2007, the NIH-funded Initiative for Maximizing Student Diversity (IMSD) program at Vanderbilt practiced GRE-free admissions (no score too low) to the first-year graduate programs [21]. This led to broadening access for PEER students to biomedical training, resulting in a sufficient number of PEER students in the IMSD program to develop a strong sense of community and the social/emotional support via peer mentorship to thrive. From 2007 to 2021, the IMSD program has supported 188 graduate students, of which 88 have completed the doctoral degree, 10 completed a master’s degree, 79 are currently in the program, 9 students did not complete the degree for various reasons and 2 are deceased. This retention rate of 89% completing or on track to complete the PhD is higher than the retention rate of non-PEER students.

The IMSD program is aligned with recommendations and best practices aimed at advancing racial equity in STEM, and intervenes at multiple levels, similar to those described by Hurtado et al. [15]. Table 1 summarizes a multi-level change strategy catalyzed by the IMSD program and implemented to address systemic barriers to racial diversity and inclusion in biomedical graduate education at the time of the mentorship education component described in this paper. These strategies were intentionally chosen because they targeted multiple levels of intervention and have a rich evidence base.

**Table 1.** Vanderbilt Model of Racial Diversity and Inclusion in the Biomedical Sciences: 2017-2021.

| Intervention level | Examples of Interventions Undertaken at Vanderbilt School of Medicine Basic Sciences (VBS) | Example Articles Demonstrating Evidence Base |
| --- | --- | --- |
| Institutional Policies, Practices, and Culture | • Replaced GRE requirement with holistic review in admissions process<br>• Instituted _BS-wide survey to assess culture and climate<br>• Initiated lecture series showcasing PEER investigators making notable discoveries as senior postdoctoral fellows/early career faculty. Provided career role models for PEER students and faculty potential candidates.<br>• Basic Science Departments and Programs established their own Diversity, Equity and Inclusion committee of faculty, staff, and students to provide programming. | • Moneta-Koehler et al. [22]<br>• Hatfield et al. [23]<br>• Hurtado et al. [24] |
| Community building | • Developed a data club and journal club for IMSD students that met regularly over dinner<br>• Hosted service-related events and outings<br>• Annual IMSD townhall meeting | • Arif et al. [25]<br>• Mishra [26] |
| Mentorship | • Careful recruitment and selection of IMSD faculty mentors<br>• Culturally Responsive Mentorship Education provided to all IMSD mentors | • Womack et al. [27]<br>• NASEM [8]<br>• Pfund et al. [14] |
| Student-level Resources and Supports | • Programming and resources related to Leadership and Management, RCR, Quantitative Skills, and Collaborative Team Science | • NASEM [8]<br>• Verderame et al. [28] |

Nonetheless, concerns remained, specifically regarding the low interest in academic careers among IMSD students who have completed or are completing the doctorate. In 2017 we (L.S., A.B.W., C.P.) launched a Graduate Student Mentoring, Climate, and Career Plans Survey to probe the training experience and career plans of Vanderbilt doctoral students across 13 different graduate programs in the biomedical sciences. Having a sample of 90+ PEER students enabled a sufficient number of responses to gather some important insights. We found that disparities existed between PEER and non-PEER doctoral students in areas such as feelings of isolation in their department or program, concerns over work life balance, and lower assessment of opportunities to discuss how their racial/ethnic or gender influences their training experience. While 30% of our biomedical doctoral students were planning to pursue an academic career, only 11% of PEER doctoral students expressed such plans. These indicators have improved and achieved greater parity over time - potentially due to some of the efforts described in this paper - nevertheless, they served to raise faculty awareness of needed change and served as a motivating factor for implementing faculty mentoring education. Certainly, there are few faculty from PEER groups, since faculty diversity in the Vanderbilt School of Medicine is considerably lower than graduate student diversity. Therefore, most PEER graduate students train with non-PEER faculty principal investigators (PIs). And while diversity has been an integral component of Vanderbilt’s biomedical graduate training programs for nearly three decades, inclusion has lagged significantly behind. The experience of belonging to the IMSD community in some sense substitutes for the lack of belonging that PEER students may experience in their mostly majority laboratory or graduate program.

It was within this climate and landscape that we (L.S., A.B.W., C.P.) introduced culturally responsive mentorship education within the Vanderbilt Basic Sciences in the late fall of 2017. The training of all doctoral students in biomedical research in the Vanderbilt School of Medicine is under the auspices of the Basic Sciences, including faculty mentors whose laboratories are in either VU or VUMC. Thus, both VU and VUMC faculty attended the mentorship education workshops. The NIH was already establishing expectations that mentorship education for faculty should be a part of T32 training grant programming and a criterion for selection of faculty to be included on T32 rosters. We took this expectation a step further to explore if improving mentorship practices could be interwoven during mentorship education workshops with dialogue on cultural awareness. Byars-Winston et al. [29], demonstrated that cultural diversity training in mentorship education increased research mentors’ awareness of the relevance of their own racial/ethnic identity to the mentoring relationship and their research mentees valued that awareness. By providing opportunities for faculty mentors to develop a stronger awareness of their own cultural identity and its relevance to effective mentorship, we posited that we could improve the research training experience for all biomedical doctoral students, and PEER students in particular. Because mentoring relationships are at the core of how mentees at all levels (graduate student, postdoctoral fellow, junior faculty) experience the research training environment, a more culturally aware faculty could in turn create a more inclusive institutional climate in the Vanderbilt School of Medicine.

### Mentorship Education Component Delivered to Vanderbilt Basic Sciences Faculty

For our mentorship education, we committed to the NASEM concept that effective mentorship is necessarily culturally responsive whereby mentors “show curiosity and concern for students’ cultural backgrounds” (pg. 62) and their non-STEM social identities [8]. We followed a mentorship education model similar to that implemented by three of the authors of this paper for the Howard Hughes Medical Institute (HHMI) Gilliam Fellowship mentors [14]. The curriculum facilitates mentorship skill development across two full days. The curriculum includes: 1) foundational mentorship education that covers specific competencies and characteristics of effective instrumental and psychosocial mentorship roles; 2) extended focus on building mentors’ cultural responsiveness to address diversity factors in their research mentoring relationships; and 3) targeted work on considering mentoring relationship dynamics within the local research training environments. The Vanderbilt mentorship education curricular elements are informed by well-studied, culturally informed mentor training interventions from the following curricula: *Entering Mentoring* [30–33], *Optimizing the Practice of Mentoring* [34], *Promoting Mentee Research Self-Efficacy* [35], and the *Culturally Aware Mentoring (CAM) Workshop* [27, 37]. Table 2 provides an overview of the number of people who participated in the Vanderbilt mentorship education workshops, workshop dates, and the formats of the workshop from 2017-2021. The content of the workshop incorporated interactive activities including small and large group discussions of readings, a short video on cultural diversity in organizations, a racial/ethnic identity self-reflection exercise, and case scenarios related to the impacts of cultural diversity on research mentee academic and career development needs and their mentoring relationships.

**Table 2.**
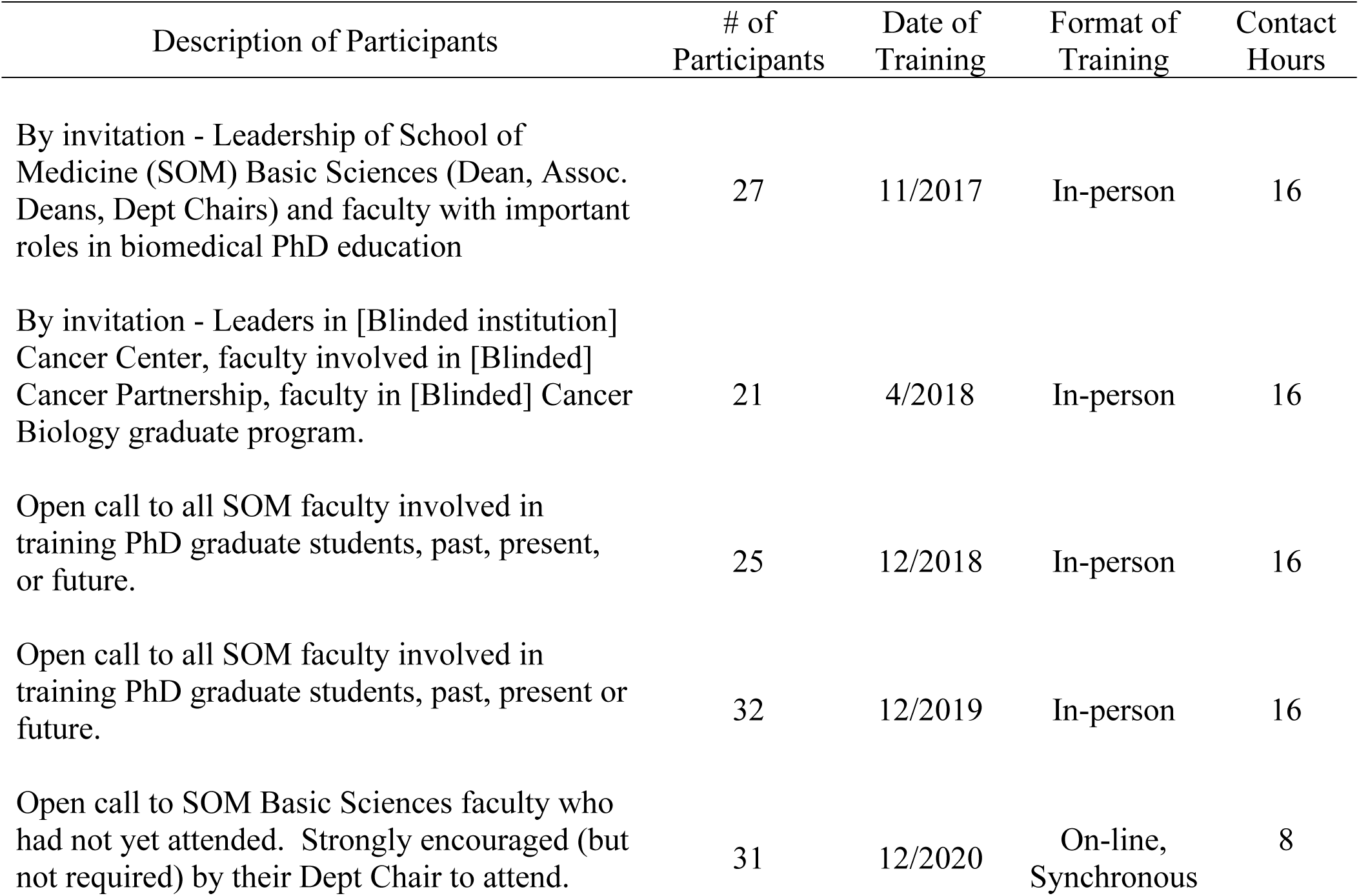
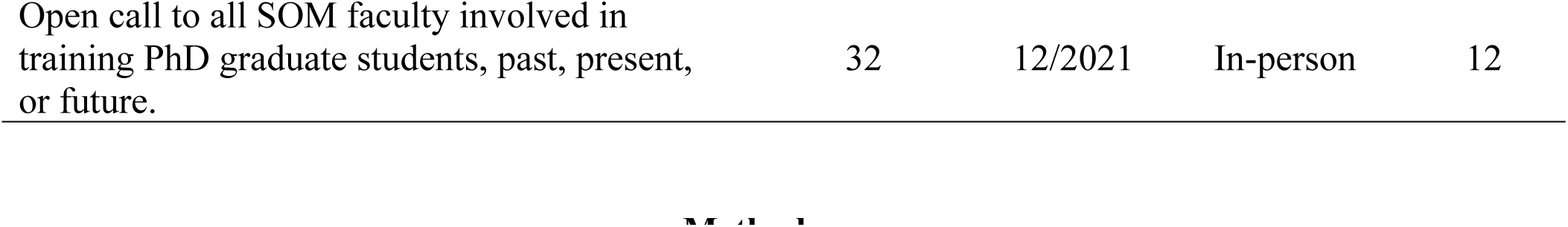
Participants in the Vanderbilt Mentorship Education Workshops 2017-2021.

| Description of Participants | # of Participants | Date of Training | Format of Training | Contact Hours |
| --- | --- | --- | --- | --- |
| By invitation - Leadership of School of Medicine (SOM) Basic Sciences (Dean, Assoc. Deans, Dept Chairs) and faculty with important roles in biomedical PhD education | 27 | 11/2017 | In-person | 16 |
| By invitation - Leaders in [Blinded institution] Cancer Center, faculty involved in [Blinded] Cancer Partnership, faculty in [Blinded] Cancer Biology graduate program. | 21 | 4/2018 | In-person | 16 |
| Open call to all SOM faculty involved in training PhD graduate students, past, present, or future. | 25 | 12/2018 | In-person | 16 |
| Open call to all SOM faculty involved in training PhD graduate students, past, present or future. | 32 | 12/2019 | In-person | 16 |
| Open call to SOM Basic Sciences faculty who had not yet attended. Strongly encouraged (but not required) by their Dept Chair to attend. | 31 | 12/2020 | On-line, Synchronous | 8 |

## Method

We used a mixed methods case study [36], and combined quantitative and qualitative data we gathered through pre- and post-surveys administered to faculty who participated in the mentorship education workshops, and a series of three focus groups involving IMSD students whose mentors had participated in the workshops. Our case study was guided by the following three research questions: 1) What types of mentoring experiences have faculty had? 2) How did faculty respond to the mentorship education? 3) How did students perceive faculty efforts at enacting culturally responsive mentoring? We sought and were granted study approval by the University of Wisconsin and Vanderbilt University Institutional Review Boards. Written informed consent was obtained from participants for the surveys and focus groups; informed consent documents were administered electronically.

### Data Collection

Data were collected through two primary methods: surveys with faculty participants, and focus groups with graduate students whose mentors had participated in mentorship education workshops.

#### Surveys with Workshop Participants

We (C.P., L.S., A.B.W.) collected data from participating mentors before the workshop (for planning and preparation purposes) and immediately following the workshop. Data were collected using Qualtrics survey platform. Questions included assessment of program components, including participant satisfaction, learning gains, mentorship quality, and outcomes, including the Cultural Diversity Awareness (CDA) measure with three subscales assessing attitudes, behaviors, and confidence to enact behaviors reflecting CDA in mentoring relationships [37].

#### Focus groups with Doctoral Students

We (S.S., L.S.) conducted three successive focus groups with students participating in the IMSD program who had been in their labs before and after their mentor participated in the mentorship workshop. Each focus group involved the same set of students, and students responded to different questions in each group. The focus groups were held on-line, via Zoom, and were spaced approximately three days apart. The first group focused on participants’ expectations regarding mentoring when they entered graduate school. In the second focus group, we asked students to reflect on their current mentoring experiences, and whether or not those experiences met their initial expectations. In the third focus group, we asked students to recommend changes that they thought would improve mentoring of students at Vanderbilt, including but not limited to PEER students.

### Participants

Because we were interested in how the mentoring experiences of students might be impacted by their mentors participating in the mentorship education training, we only asked IMSD students to participate in the focus groups if they were being mentored in a lab before and after their mentor participated in the training. All faculty who participated in the mentorship education workshops were invited to take the pre- and post-surveys. Of the 168 workshop participants, 117 completed surveys and consented to their use for workshop planning purposes; 108 completed the surveys and consented to their use for research purposes, yielding a survey response rate of 64% for faculty returning both pre- and post-surveys.

#### Faculty Participants

Demographic questions as well as questions related to participants’ reasons for participating in the training were included in our study pre-surveys. Table 3 provides demographic descriptors of faculty participants by year. Participants varied across rank and level of leadership, with the highest percentage of participants across all years being Full Professors (40%), followed by Assistant Professors (30%), and Associate Professors (19%). Twelve percent of respondents indicated that they held a leadership position, either as an Assistant Dean, Associate Dean, or Dean (7%) or as a Training Program Director (5%). The majority of participants identified as White (73%) with very few faculty identifying with a PEER group (8% reported American Indian or Alaskan Native, Black or African American, Native Hawaiian or Pacific Islander; 6% reported Hispanic/Latino). Ten percent elected not to report their race; 16% did not report their ethnicity. Forty-seven percent of participants identified as men, 43% of participants identified as women, and 10% of participants did not report their gender.

**Table 3.** Sociodemographic Characteristics of Workshop Participants.

| Sample Characteristics | % |
| --- | --- |
| Gender |  |
| Man | 47 |
| Woman | 43 |
| Did Not Report | 10 |
| Race |  |
| White | 73 |
| Asian | 9 |
| Black or African American | 6 |
| American Indian or Alaskan Native | 1 |
| Native Hawaiian or Pacific Islander | 1 |
| Other race not listed | 3 |
| Did not report | 10 |
| Ethnicity |  |
| Hispanic/Latino | 6 |
| Not Hispanic/Latino | 78 |
| Did Not Report | 16 |
| Title |  |
| Assistant Professor | 30 |
| Associate Professor | 19 |
| Professor | 40 |
| Assistant Dean, Associate Dean, or Dean | 7 |
| Training Program Director | 5 |
| Other Title Not Listed | 7 |
| Did Not Report | 9 |

*Note* N=108; Participants could select more than one race and more than one title

#### Student Participants

When the focus groups were conducted, first and second year students were excluded from the study because of the impact of the COVID-19 pandemic on their early graduate school experiences. Specifically, we believe their experiences with mentoring, course work, and lab work were meaningfully different enough from previous students to justify their exclusion from this study. The number of students who met study criteria for each year in a Vanderbilt Basic Sciences doctoral program are as follows: 2015 (6th year students), n = 3; 2016 (5th year students), n = 15; 2017 (4th year students), n = 6; 2018 (3rd year students), n = 2. We invited all of these students to participate with the goal of recruiting 6 to 10 participants [38] and yielded 8. Student participation was spread across years of the program, with the most students coming from years 4 and 5. We do not report participant race and gender in an effort to protect the confidentiality of those who participated, although all students were IMSD program participants, and thus come from PEER groups.

### Data Analysis

We (F.S., C.P.) focused our analyses of the survey data on faculty participants’ values related to mentoring, mentoring confidence, and use of specific mentoring behaviors, as well as their motivations for participating in the workshop and their satisfaction with the mentoring education they received. These data were analyzed for all six faculty participant cohorts 2017-2021. Our analysis of mentee focus group data explores mentees’ perceptions of the mentoring they received at Vanderbilt, while also examining the broader context and IMSD-driven changes mentees experienced.

#### Quantitative analysis

Post training survey data from across cohorts were cleaned and combined using Microsoft Excel. Population means and standard error from across cohorts were calculated using Excel. Boxplots were generated using the ggplot2 (https://ggplot2.tidyverse.org/) R package.

#### Qualitative analysis

Qualitative data from open-ended survey questions were analyzed using an iterative coding methodology [39]. For the first round of coding, open coding was employed by co-author C.P., followed by some code consolidation and suggestions by co-author F.S. Focus group data were analyzed by co-author S.S. using a grounded theory approach [40] that involves reading transcripts and assigning codes to words or phrases in the transcripts and then exploring the prevalence and relationships among codes to develop concepts and theories from the data. When reporting qualitative data, we used participant identification numbers rather than names or demographic descriptors to maintain the confidentiality of the students. Additionally, we have used the gender-neutral pronoun “they” (versus “he” or “she”) when describing students to further conceal their identities. There are times when participant responses reveal their gender.

When these revelations are important to understanding the participant quotation, we have left them in. When students are discussing their mentors, we have retained demographic descriptions when those descriptions are important to understanding the data.

## Results

### Faculty Experiences with Mentoring and Mentorship Education

Faculty participants had varied experience with mentoring graduate students, ranging from faculty who had never mentored a graduate student (3%) to faculty who had mentored more than 11 graduate students (35%). On the whole, workshop participants had extensive experience with mentoring, with 62% indicating that they had mentored 6 or more graduate students. These trends were reversed when faculty were asked about their experience mentoring graduate students from PEER groups. Fifty-nine percent of faculty reported that they had mentored 2 or fewer students from PEER groups, with only 6% reporting having mentored 11 or more. Despite the vast majority of participants having mentored at least 1 graduate student, only 42% of faculty reported having previously participated in mentor training of any kind.

### Faculty Motivation for Participating in Mentorship Education

Of the 168 participants who completed mentorship education workshops, 94 (57%) responded to an open-ended prompt on the pre-survey that inquired, “Why did you sign up for the workshop?” The responses revealed a number of motivations, such as desire to improve ability to mentor trainees from diverse backgrounds (40%), desire to improve mentoring skills generally (27%), being invited or strongly encouraged to attend (12%), and preparing for a new campus role (4%). Although most participants appeared to have taken part in the workshops because they were motivated to improve their mentoring in one way or another, some participants were more reluctant. For example, one participant indicated they attended because, “my university now considers having such training as necessary to allow PIs to mentor PEER students.” Another said they attended because of “influence from a local colleague invested in such matters.” Table 4 displays themes in participant responses around motivation for participating, as well as counts and sample quotations.

**Table 4.**
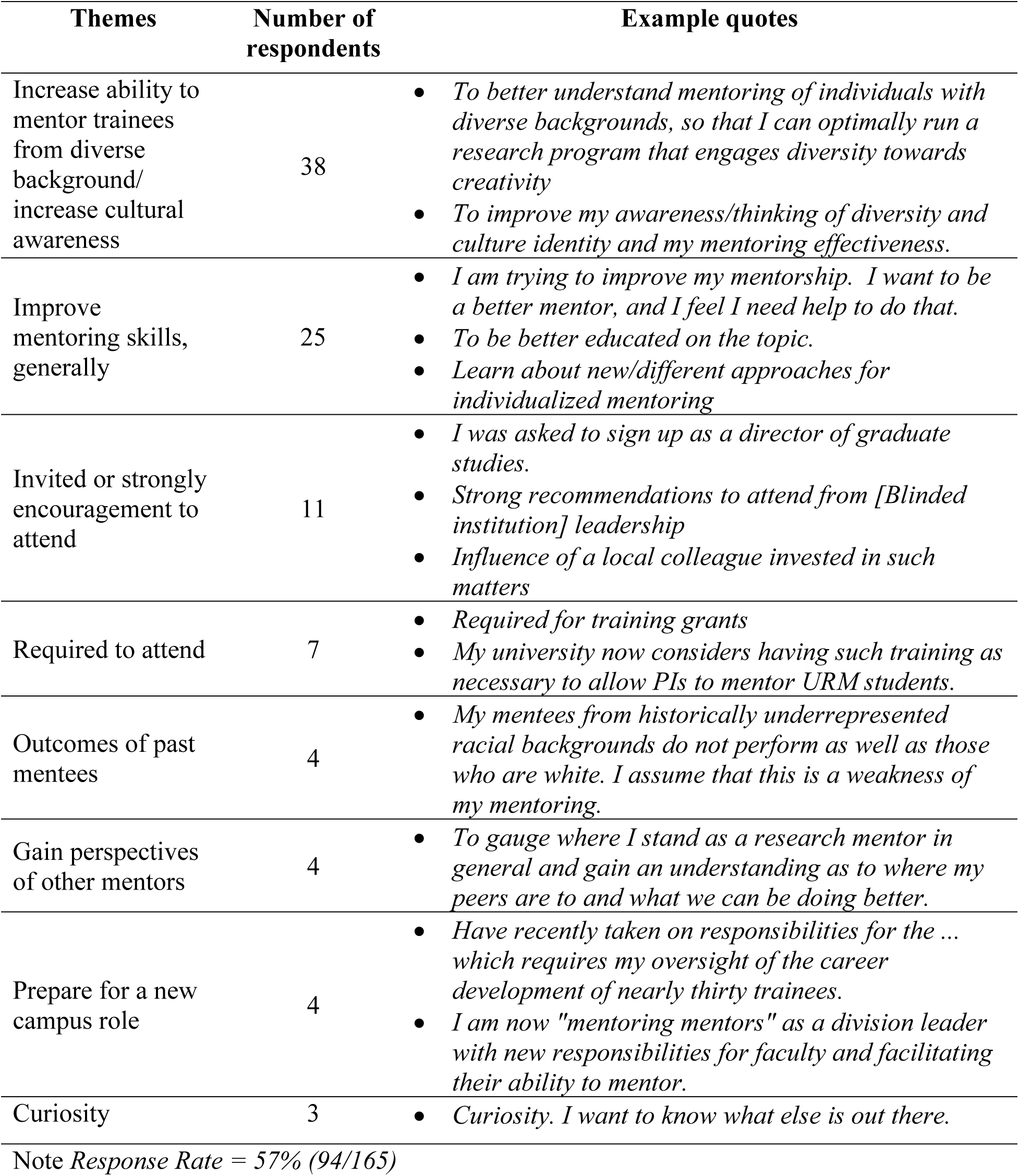
Faculty Motivation for Participating in the Workshop.

### Faculty Perceptions of Their Mentoring Effectiveness

In response to items intended to assess participant perspectives on their own mentoring effectiveness and quality following mentorship education, faculty rated their mentoring skills favorably across participant cohorts. Figure 1A demonstrates response distribution (Likert-type scale 1 [strongly disagree] to 4 [strongly agree]) across all six cohorts to questions regarding mentoring effectiveness including the mentor’s showing interest in student projects (n = 105), and their efforts to make students feel included in their lab and lab culture (n = 104). Although there are some faculty who responded “strongly disagree (1)” or “disagree (2)” in a few of the participant cohorts, on average, faculty rated themselves positively on both (3.88 ± 0.47 for showing interest; 3.62 ± 0.61 for inclusion). Similarly, faculty rated themselves highly on items intended to measure perceptions of mentoring quality. Across the participant cohorts, the majority of faculty rated their working relationship with their mentees as “good (3)” or “excellent (4)” (3.38 ± 0.52), and the overall quality of their mentoring relationships “good” or “excellent” as well (3.23 ± 0.56). Figure 1B displays response distribution across all six cohorts.

**Figure 1.**
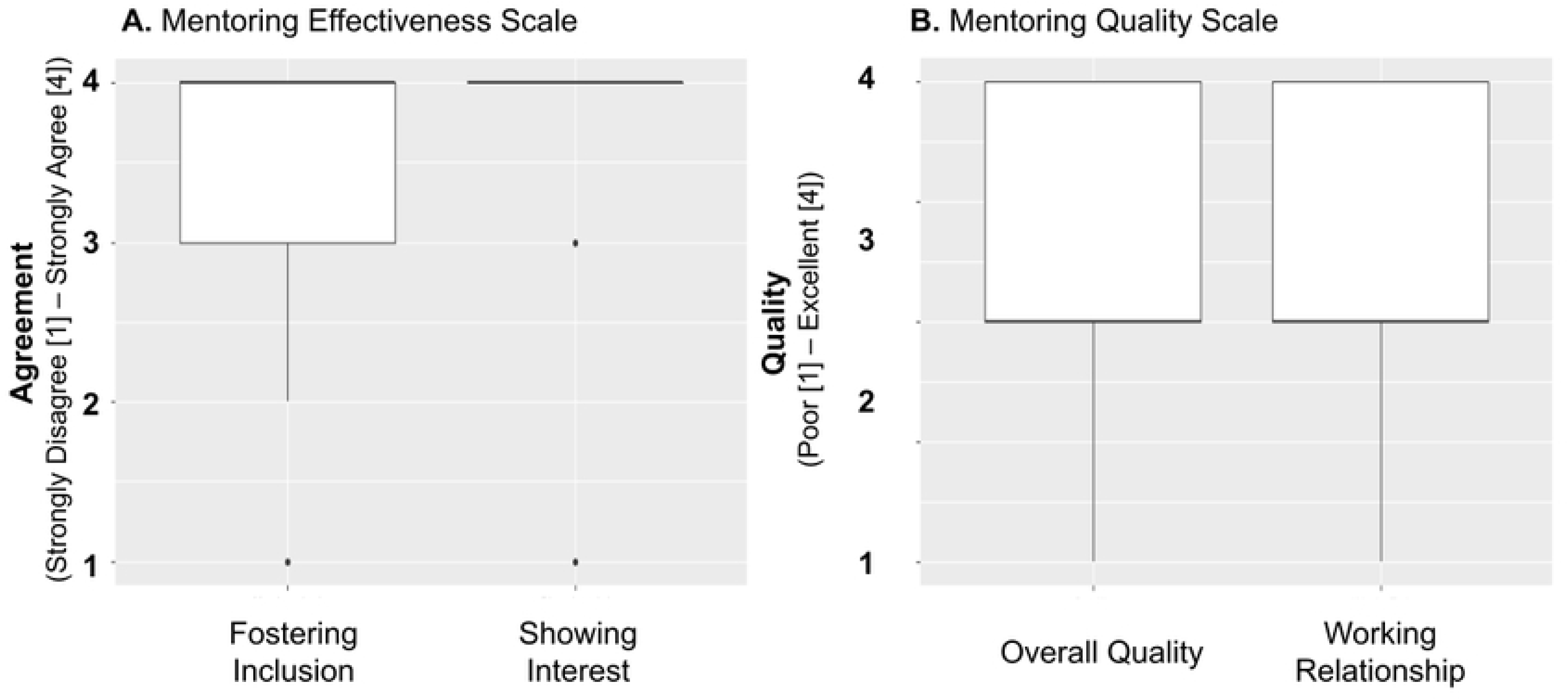
Self-Reported Mentoring Effectiveness and Quality. A. Self-reported mentoring effectiveness following training for all six cohorts. Mentors rated their agreement using a Likert-type scale from 1 (strongly disagree) to 4 (strongly agree) on the following items: “I made my mentee feel included in the lab (or field setting)” and “I tried to show interest in my mentee’s projects.” B. Self-reported mentoring quality following training’ for all six cohorts. Mentors rated the quality of two aspects of their mentoring relationship (“The overall quality of my research mentorship relationship”; “My working relationship with my research mentees”) using a Likert-type scale ranging from 1 (poor) to 4 (excellent).

#### Culturally Diversity Awareness (CDA) in Mentorship: Attitudes, Confidence, and Behaviors

Similar to previous implementations of culturally responsive mentorship education [14, 29] faculty reported high levels of agreement with CDA Attitudes scale items such as, “*It is important for mentors and mentees to talk together about the mentee’s racial/ethnic background*,” and “*It is important for mentors and mentees to discuss how race/ethnicity impacts the mentee’s research experience.*” Figure 2A displays response distribution across all six cohorts for CDA Attitudes, as well as CDA Confidence and Behaviors (discussed below). We have collapsed data across all cohorts for simplicity in viewing and interpreting figures, and because a Kruskal-Wallis test revealed that cohorts were statistically indistinguishable from one another when comparing means and distributions of all items, save two: “My racial/ethnic identity is relevant to my research mentoring relationships,” (Attitude item, p = 0.032) and “I intentionally created opportunities for my mentees to bring up issues of race/ethnicity as they arose” (Behavior item, p = 0.024).

**Figure 2.**
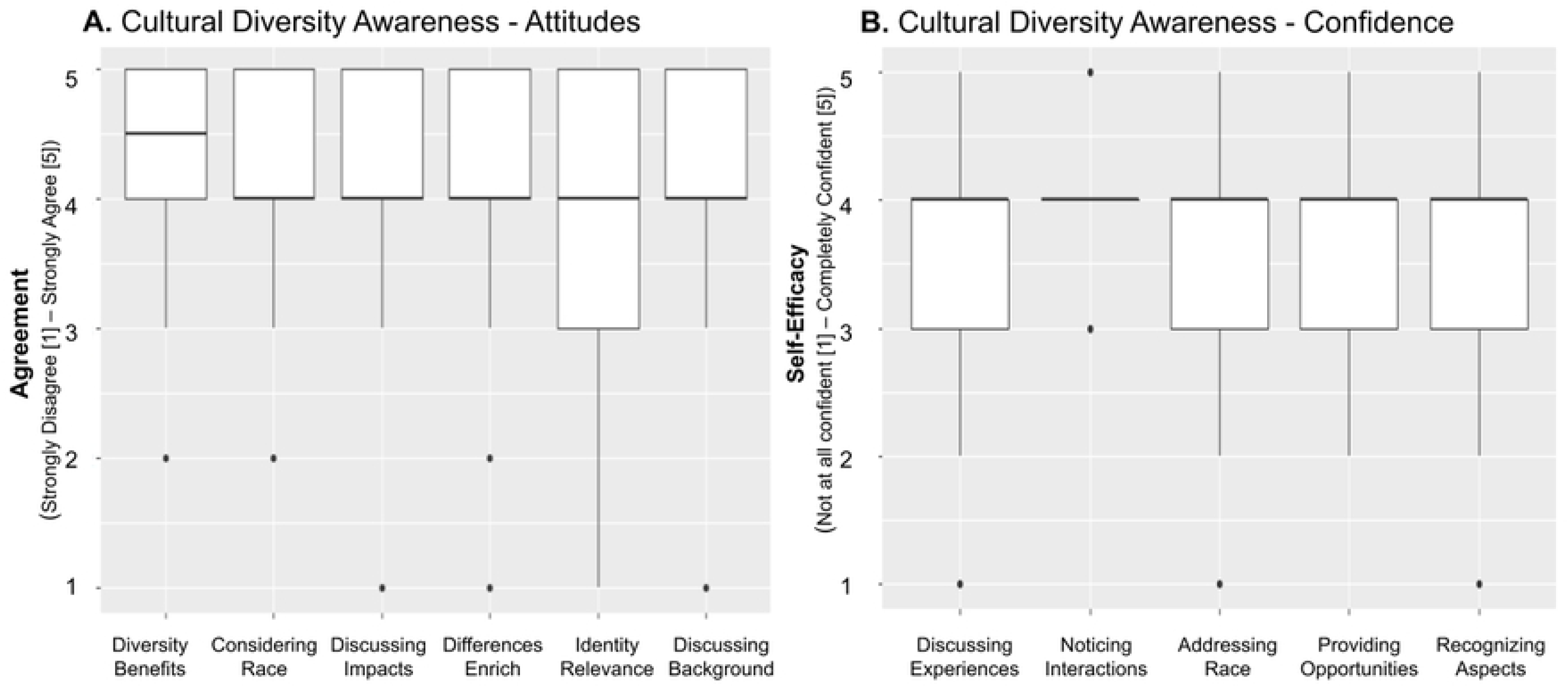
Cultural Diversity Awareness and Confidence. A. Self-reported attitudes towards cultural diversity awareness (CDA) from all six cohorts following training. Respondents rated their agreement on a Likert-type scale ranging from 1 (strongly disagree) to 5 (strongly agree) to the following statements: “Mentoring someone with a different racial/ethnic background benefits the research”; “It is important to consider the mentee’s and mentor’s race/ethnicity in mentoring relationships”; “It is important for mentors and mentees to discuss how race/ethnicity impacts the mentee’s research experience”; “Racial/ethnic differences between mentors and mentees enriches the research mentoring group”; “My racial/ethnic identity is relevant to my research mentoring relationships”; “It is important for mentors and mentees to talk together about the mentee’s racial/ethnic background” B. Self-reported confidence to enact various behaviors reflecting CDA from all six cohorts following training. Respondents rated their confidence on a Likert-type scale ranging from 1 (not at all confident) to 5 (completely confident) on the following skills: “Discuss with mentees how it feels to be a minority in science”; “Notice interactions in the mentoring relationship that could be insulting or dismissive of mentees because of their race/ethnicity”; “Take advantage of opportunities to address race/ethnicity in the research mentoring relationship”; “Provide opportunities for mentees to talk about their racial/ethnic identity as it relates to their research experience should the occasion arise”; “Recognize aspects of the research mentoring experience that may make racial/ethnic minority students feel vulnerable to confirming stereotypes.”

**Figure 3.**
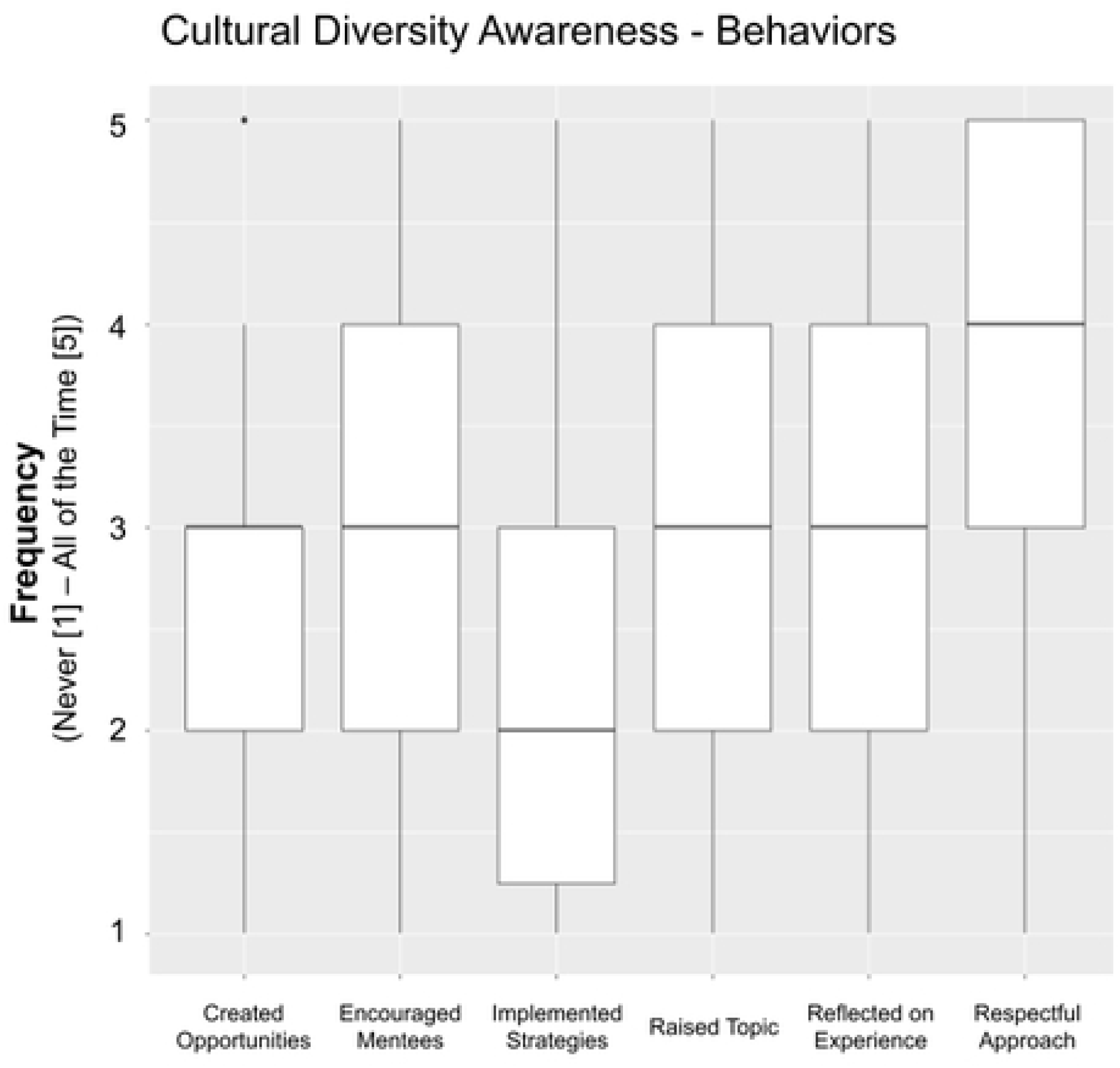
Self-reported frequency of culturally aware mentorship behaviors. Mentors from all cohorts were asked to rate the frequency with which behaviors occurred in their mentoring relationships in the last year. Answers were on a Likert-type scale ranging from 1 (never) to 5 (all of the time). Behaviors included the following: “I intentionally created opportunities for my mentees to bring up issues of race/ethnicity as they arose”; “I encouraged mentees to talk about how the research relates to their own lived experience”; “I implemented specific strategies to address racial/ethnic diversity in my research mentoring relationship”; “I raised the topic of race/ethnicity in my research mentoring relationships when it was relevant”; “I reflected on how the research experience may differ for mentees from different racial/ethnic groups”; “I approached the topic of race/ethnicity with my mentee(s) in a respectful manner.”

Faculty ratings of their confidence to enact CDA behaviors were comparatively lower in absolute value than ratings of their attitudes toward the relevance of CDA in mentorship. This is true for all questions and all cohorts, with the exception of the item “Notice interactions in the mentoring relationship that could be insulting or dismissive to mentees because of their race/ethnicity,” for which most faculty rated themselves as confident or completely confident.

When asked a follow up question about their confidence in mentoring someone of a different race and ethnicity, however, they reported high levels of confidence with an average of 5.40 ± 0.98 on a 1 to 7 scale, where 1 denotes “very low” confidence and 7 “very high” confidence.

When asked to report the degree to which they implemented culturally responsive mentoring behaviors in the past 12 months, participants recorded answers ranging from “Never (1)” to “All of the time (5),” though the majority of participants responded somewhere between the two extremes. Among the items, “I approached the topic of race/ethnicity with my mentee(s) in a respectful manner” had the highest percentage (64.4%) of survey respondents indicating that they did this “frequently,” or “all the time.” The item “I implemented specific strategies to address racial/ethnic diversity in my research mentoring relationship,” had the lowest percentage of faculty (21.6%) indicating that they did this “frequently” or “all of the time.”

Bivariate correlations between the three CDA subscales revealed significantly positive relationships among the variables: Attitudes and Behaviors (r =.33); Attitudes and Confidence (r =.41); Behaviors and Confidence (r =.44). The small to medium effect sizes indicate that although the variables are interrelated, there are unique aspects of how faculty mentors perceive and engage in CDA in their mentoring relationships.

#### Workshop Satisfaction and Outcomes

After the workshop, participants were asked to rate their satisfaction with aspects of the workshop. In these surveys, 93% indicated that the materials were useful, and 95% that the workshop was well-organized and easy to follow. Ninety-nine percent agreed that the speakers were knowledgeable, and 99% indicated that the workshop was delivered in a respectful and sensitive manner. The lowest satisfaction rating had to do with time, with 18% of participants indicating that there was insufficient time allocated to the workshop. Participants also reported high satisfaction with the different modules covered in the training. These ratings did not demonstrate significant differences among modules or cohorts. Perhaps most importantly, 97% participants indicated that they were likely or very likely to make changes to their current mentoring practices based on something they learned at the workshop. Similarly, 98% indicated that they would recommend the workshop to a colleague. Based on the popularity of the workshop over time, it appears that they did - later years were full prior to being able to accommodate everyone who was interested in participating.

### Student Perceptions of Mentoring

Faculty intending to implement culturally responsive mentoring practices is a critical component of improving mentorship, but understanding if these behaviors are implemented and how they are perceived by mentees is the true test of change. We conducted focus groups with 8 3rd – 6th year IMSD students whose mentors had attended the mentorship education workshop during the time the student was working in their lab to garner some of these perceptions. Overall, the students we interviewed from the IMSD program were happy with the mentoring they were receiving at Vanderbilt. They mentioned previous positive mentorship experiences (e.g., in undergraduate programs) and the ability to change mentors once at Vanderbilt as experiences and opportunities that helped them identify what they wanted in a mentor and develop strategies for receiving it. Students also noted being satisfied with diversity among doctoral students at Vanderbilt, but disappointed in the lack of faculty diversity. One student noted,

> One of the nice things about the IMSD program is that you know 30% of your peers will be people of color, so you know there is diversity coming in. But there isn’t much diversity in the faculty…it is challenging to go into academia…or industry or wherever…when you don’t see faculty and leadership that looks like you. (Focus group 1, Participant 2)

Alongside lack of faculty diversity, students noted varied willingness and capabilities among faculty when it came to mentoring PEER students, and using strategies known to be aligned with culturally responsive mentoring. Students’ responses about their mentors centered on themes related to faculty creating opportunities to discuss race, helping students connect research with lived experience, and observations about institutional structures and policies that incentivize good mentoring.

#### Creating Opportunities to Discuss Race

When students were asked if their mentors were comfortable with and willing to bring up race, most students reported that they were. That said, students emphasized that *how* mentors brought up race and talked about it mattered just as much as their willingness to do so.

> So, my mentor also talks about race quite a bit…almost every conversation that we have, it comes up, especially within the past year. And he’s somebody who he feels like in every aspect of life, his duty is to mentor and he wants to give advice…he just feels like he needs to give me advice all the time, and the race thing is not one where I want to talk to him about it because he doesn’t know anything about it, and I think he wants to understand and he’s trying to be empathetic, but I have no interest to talk to him about that at all. So I kind of resent every time he brings it up. (Focus group 2, Participant 3)

This viewpoint was reinforced by another participant.

> Yeah, I’ve had that same exact experience with my mentor. Whenever a topic of race or diversity would come up, he would love to talk about how he’s first generation and so that makes him diverse, and basically try to equate our situations - him as a white man and me as a black woman are the same thing because he was first gen, and I’m like that’s not the same thing. (Focus group 2, Participant 4)

Other students spoke up and contrasted these experiences with mentoring that they identified as more positive. A difference seemed to be that mentors whose willingness to raise issues of race was perceived as helpful when people raised them in the spirit of making space for discussion rather than sharing opinions or comparisons, as the following exchange demonstrates:

> My mentor is one of those individuals where she’s aware of these things that are going on, like one particular example is when the George Floyd verdict came out, she understood (the significance) of what was going on and asked, “how are you feeling about everything?” She allows me to be open with her and have that open conversation, so I definitely think she’s pretty good at having those certain conversations. I guess she identifies as a white woman and she knows she only knows things to a certain extent…but I think that she’s willing to have an open ear and to listen and, you know - I’m here for you if you need anything…I feel okay having those sorts of conversations with her - it never feels too awkward. (Focus group 2, Participant 6)

Similarly, another student said,

> Yeah, my mentor is also open to talking about stuff to do with race, we actually - last year with when the protests happened - we had a lab conversation about race and she made it very clear that if we ever want to talk about something with her or we feel like we need to take a day off to deal with what’s going on in the news, or whatever that we can do that. So it’s not brought up very often but she made it very clear that it’s okay to talk about and I think everyone goes comfortable talking about that stuff with her. (Focus group 2, Participant 8)

#### Connecting Research with Lived Experience

In terms of mentors helping students connect their science to their personal lives, an important aspect was acknowledging the validity and value of educational and professional goals that were different than their mentor might have pursued. These goals included living in a particular area of the country to be close to family members (even if that limited a student’s job prospects), and being responsive to student desires to work outside of academia or an R1 university environment. Describing a positive experience with their mentor, one student said,

> So, for example, I was writing a grant, and as part of that we have our career statements. And I’ve always said one of the things I want to do when I finish my PhD is to go back and teach at an historically black college, because that’s where I went to school. My mentor has never downplayed that…she’s never pushed me to go R1. She’s always like ‘okay, you know the stuff that you want to do.’ This has allowed me - as much as I want to - it allows me to kind of bring myself into my work and my goals. (Focus group 2, Participant 6)

This theme emerged again in the later focus group about what a person would want from an ideal mentoring relationship; how someone would want to change their experience with mentoring:

> I think the first thing would be to listen to what I want about my future goals more. So I think my PI, kind of just assumes everyone wants to be a PI at an R1 like he is, and he doesn’t really change his training based off of his trainees goals. (Focus group 3, Participant 4)

> I could add to that, so aside from what was just said, is letting people communicate as to what they want for their future goals. In my experience - and talking to postdoc labs and interviewing (for jobs) there are people who rank their trainees - and the ones that are the best they’re like ‘oh you should be a faculty, and everybody else can do other things.’ So just being more open to other options, as opposed to putting it onto people what they should do. (Focus group 3, Participant 1)

In addition to wanting mentors who recognized and appreciated the validity of different professional goals, students appreciated when mentors encouraged them to bring their “whole selves” into their work. Although some students were clear that it was important to them for their personal and professional lives to be separate, other students wanted mentors who knew them personally and were willing to work with them around their individual strengths, limitations, and passions. In an exchange between three students, participants noted the importance of having mentors who would work with them when they needed extra support, time off, or flexibility in terms of work requirements:

> PI has been pretty amazing, as to letting me do things when I can do them…She’s been remarkably flexible with what’s going on in my own life and allowing me to work through my PhD on my own terms, and my committee has been pretty fantastic about that too. And so, even though I’ve been sick. It hasn’t like slowed down my progress, so that’s been great. (Participant 5)

> I’ve had a similar experience. (Participant 7)

> Yeah, I had a personal issue last fall, and I felt comfortable going to my mentor talking about it, and I was just in a really bad place, and I didn’t go into work for a week and she was fine with that. And, and I felt really supported, knowing that, and she’s just really supportive in general. (Participant 8)

For other students, bringing one’s “whole self” meant having a mentor who recognized their strengths and passions. One student described how her mentor gave her a platform for pursuing her passion for supporting women and minorities in science. She said,

> I would say also a really good mentor will see someone’s passion in science or even outside of science. So I have a lot of passion for minorities and women in science and my PI could definitely see that. And when the documentary Picture a Scientist came out, which talks about harassment against women in science, he asked me to co-facilitate a lab meeting about the documentary with my lab, which I was so excited about. And I was just very happy that he was able to see the amount of passion I had, and to want to work together to show that as a PI he is standing with a female in science and being like, ‘Hey lab, let’s talk about this and learn about how we can all improve in this field.’ I think a mentor that can see your passion for anything - even if it’s not data driven at the bench doing experiments - helps to see the entire scientist as a human being with passions and desires that are science oriented but also can move outside of the very structured, you know, data driven numbers that we’re used to seeing. (Focus group 3, Participant 1)

#### Institutional Incentives for Mentoring

At the end of the focus groups, we asked if students had anything else they wanted to share with us about mentoring – questions we didn’t ask but should have, other things that had occurred to them along the way, etc. This opened up an interesting and insightful conversation about how the university incentivizes mentoring and discourages racism among faculty. What was most important about this conversation is the insight the students had about problems of mentoring and discrimination of various types (racism, sexism) being structural problems within the university as opposed to strictly being about shortcomings of individual people and their ability (or lack thereof) to mentor under-represented students. For example, students suggested that the University and/or Department should collect data on student experiences with particular mentors, and use that information when students are choosing faculty mentors.

> I don’t know if it’s possible to do this, but it seems like it would be good to look at the history of the PI as a mentor before pairing (them with students)…I mean, I know we pick our mentors, but if there was a committee that looked at the relationship between mentors and especially diverse students to avoid problems in the future…I don’t know. It seems like that would be beneficial. (Participant 4)

Another, similar suggestion had to do with working with faculty in a more targeted way to improve their mentoring:

> I guess also, going off of what others are saying in terms of feedback for professors. I currently don’t know if there is a mechanism in place where mentors get feedback - I don’t know if it would be from the chair of the department or the dean – but in terms of if they do see patterns. (For example, the chair or dean saying to the faculty member) ‘You’ve had four people leave your lab in the last three years. Are there issues that we need to address or that we can change?’ and then provide targeted training or something like that. I’m just trying to think of how to make the situation better so that a mentor doesn’t get worse and worse…So like with students where we have these individual development plans that we sit down with our mentors and go through - Is there an IDP for the faculty? Where do they get their feedback, and also how important do they even see that feedback if it’s not research driven? I know a lot of the R1 faculty say that ‘research is our top priority. Everything else comes secondary. If it’s not for a grant or for a paper, I don’t have time for it.’ So another question is how do we incentivize or change the perspective (of the mentors) so that the students who are creating this data for your grants and publications, they need to have high quality mentorship from you as well. (Focus group 3, Participant 3)

## Discussion

Mentorship education interwoven with dialogue on cultural awareness was introduced by the Vanderbilt Basic Sciences to improve the research training environment with the long-term goals of improving climate, sense of belonging, and interest in staying in academia for doctoral students from PEER groups. In this mixed methods case study, we found evidence for the positive impact of faculty participation in this mentorship education on both the perceptions of mentors, who predominantly self-identified as white, and their PEER mentees, consistent with previous research [8, 14, 32]. The results of our study demonstrated that not only were mentors satisfied with the mentorship education, but the education also had a positive impact on mentors’ self-ratings of the quality of their mentoring relationships and their mentoring effectiveness.

Overall, the PEER doctoral mentees of these faculty mentors noticed and valued the improved efforts of their mentors to be culturally aware in their mentoring relationships. We highlight several findings and their implications for systemic efforts to diversify STEM, referencing components of the Hurtado et al. [15] organizational learning (OL) model for advancing inclusive science where relevant.

First, our findings reveal that faculty have varying motivations for participating in mentorship education, especially focused on increasing their cultural awareness in mentorship. Similar to research from Butz et al. [14], although many faculty in our study were intrinsically motivated to participate in mentorship education, some did so because of their faculty colleagues’ encouragement. This highlights the role of positive peer influence in increasing faculty buy-in to engage in mentorship education. Byars-Winston et al. [42], found that culturally aware-focused mentorship education promoted behavioral changes not only in faculty mentoring practices, but also in their relationships with faculty colleagues and administrators. Following Hurtado et al.’s [15] OL model, mentorship education is an important factor to create faculty buy-in to gain new knowledge about PEER trainees that change faculty mindsets and mentorship behaviors.

Second, our findings revealed a value-action/will-skill gap for faculty mentors in their cultural diversity awareness. Specifically, where many reported high endorsement of the importance of culturally diversity awareness (CDA) in their mentorship, they reported comparatively lower confidence to enact behaviors consistent with that awareness. This finding is consistent with House et al. [43], who also found that faculty mentors reported challenges in their self-efficacy to enact actual culturally aware mentoring behaviors. Findings from our two-day workshop can raise faculty awareness and provide strategies for advancing culturally responsive mentorship practices. But institutions would do well to leverage such interventions to build opportunities for continued learning and practice, consistent with Hurtado et al.’s [15] OL model for changing faculty mindsets and behaviors. Ongoing mentorship skill-building can be supported by creating opportunities for faculty to share and disseminate knowledge and skills related to effective mentorship, like communities of practice.

Third, student responses demonstrated that faculty were implementing culturally responsive mentoring practices, and – for the most part – they were appreciative of these efforts. Students noted that some faculty efforts landed better than others, and that some mentors were more skilled in culturally responsive mentoring practices than other mentors. For example, most students noted that their mentors made space to discuss race when relevant and listened to the students’ experiences. The students were also clear that *how* mentors broach the subject of race was critical to whether it is helpful or hurtful. When non-PEER faculty raised the topic of race and approached it from a position of expertise, this was received as off-putting by students.

Although it goes without saying that a white mentor conveying expertise about race to a PEER student is likely to be unhelpful, this points to an expertise paradox for some faculty. Whereas faculty mentors hold scientific expertise – and thus are rightly leaders in that realm, students hold expertise about their life experiences and needs that should be regarded as equally important in their professional training. Nathan, Koedinger, and Alibali proposed the concept of expert blind spot to capture the overgeneralization of one’s expertise in a given to another domain [44].

Relative to culturally responsive mentorship, the experiential expertise of PEER students may eclipse the disciplinary expertise of faculty mentors, thus disrupting well established hierarchies with respect to professional rank (which is earned) and race (which is not). This disruption necessitates that faculty share power with their students instead of power over them.

The notion of cultural humility may be useful to faculty mentors enacting culturally responsive mentorship practices. Cultural humility [45] is an interpersonal stance that is other-oriented rather than self-focused and involves a willingness to reflect on oneself as a cultural being and openness to hearing and understanding others’ cultural backgrounds and social identities. Thus, culturally humble faculty mentors display a respectful curiosity for others’cultural identities, resist the human tendency to view their personal beliefs and values as superior, are mindful that their understanding of others’ cultural backgrounds is limited, promote a culturally self-aware mindset, and view the practice of cultural humility as a lifelong learning process [46].

Finally, students recommended that Vanderbilt as an institution develop policies that relay the value of mentorship. This recommendation came without specific prompting, mostly offered in response to the final question of the focus group, “What else do you want me to know?”. This is noteworthy considering that mentorship is often considered a practice that operates and creates change at the interpersonal level. Students, however, seemed to see the need for institutional-level policies to ensure effective mentorship, recognizing a significant opportunity for improved mentorship to lead to shifts in the institutional climate. The NASEM report, *The Science of Effective Mentorship in STEMM* (2019), presents several recommendations university leaders can embrace to advance a culture of mentorship at their institutions. These include:

- Develop a shared vision of goals for degree attainment, job placement, and professional development in STEMM that includes mentorship as a component.
- Engage faculty professional development programs and centers in addressing mentorship as part of undergraduate research, graduate training, faculty learning communities, new faculty orientation, and regular programming.
- Provide funding to facilitate mentor-mentee activity surrounding students’ research interests.
- Encourage campus-wide promotion review committees to establish guidelines for evaluating mentorship activities and impact.
- Encourage campus STEMM programs and other student success programs to evaluate and report on key mentorship components when reviewing overall program impact.

The students’ recommendation emphasizes the positive impact of social norming on faculty behaviors that departments and training programs can communicate: “formal mentorship education and effective mentorship are valued and expected here.” Indeed, Vanderbilt Basic Sciences has made some progress in this regard, now requiring faculty to participate in an in-person mentorship education session in order to accept doctoral students into their laboratory.

Changing social norms is a critical part in Hurtado et al.’s [15] OL model. The theory of planned behavior [47] suggests that injunctive norms, or the expected behaviors in a given social context - like a department - will become reinforced in that context. Change the institutional expectations, and people will feel more compelled to follow them.

### Limitations

Although we complied with best practices in case study methods such as employing a well-bounded case, triangulating data sources and data types, and working iteratively through data analysis, interpretation, and author reflection, there are nevertheless limitations. Survey response rate was 64% across all cohorts for survey data, and 8 out of 26 invited students elected to participate in the series of three focus groups. We do not know whether the non-responses reflect lack of time, lack of interest, or something else; nor do we know whether non-responders would have shared similar perspectives to those who participated. Furthermore, because our case study does not follow individuals across time, we did not assess individual change with respect to mentoring behaviors or the durability of mentors’ intentions to change. Finally, we did not conduct within-group data analyses nor use an intersectional lens with respect to mentor or student experiences. These experiences *are* represented in the qualitative data to some extent, however, we did not probe for them. As a result, we have missed some of the ways in which gender, class, sexuality, and other identity categories intersect with race to shape mentor-mentee relationships and their potential for driving institutional change.

### Conclusion

There are promising approaches to helping faculty becoming more culturally aware and to improve their skills in culturally aware mentoring practices [14, 20, 29, 42, 43]. Findings from this present study add to the call for research training environments to institutionalize culturally responsive mentorship efforts [8].

## Acknowledgments

In addition to grant funding, this study was supported by the Office of the Dean of Basic Sciences, Vanderbilt School of Medicine; the Center for the Improvement of Mentored Experiences in Research (CIMER) in the Wisconsin Center for Education Research at the University of Wisconsin-Madison; and the Department of Medicine at the University of Wisconsin-Madison. We would like to acknowledge the staff of the Vanderbilt Office of the Dean of Basic Sciences, especially Tracy O’Brien, Becky Sanders, Steven Doster, and Anne Lara, for their assistance with workshops, and the unwavering support of Larry Marnett, Vanderbilt Basic Sciences (VBS) Dean Emeritus and Jennifer Pietenpol, Chief Scientific and Strategy Officer, VUMC. We thank the Vanderbilt Office of Biomedical Research Education and Training (BRET), in particular Roger Chalkley and Abigail Brown, for assistance with the doctoral student Climate, Culture and Career Plans survey, as well as Felysha Jenkins, VBS Assistant Dean for Diversity, Equity and Inclusion and Larry Marnett for helpful comments on the manuscript. Finally, we thank the CIMER Evaluation and Research team for their assistance with mentorship education evaluation data collection.

